# HumanMine: advanced data searching, analysis and cross-species comparison

**DOI:** 10.1101/2022.05.11.491545

**Authors:** Rachel Lyne, Adrián Bazaga, Daniela Butano, Sergio Contrino, Joshua Heimbach, Fengyuan Hu, Alexis Kalderimis, Mike Lyne, Kevin Reierskog, Radek Stepan, Julie Sullivan, Archie Wise, Yo Yehudi, Gos Micklem

## Abstract

HumanMine (www.humanmine.org) is an integrated database of human genomics and proteomics data that provides a powerful interface to support sophisticated exploration and analysis of data compiled from experimental, computational and curated data sources. Built using the InterMine data integration platform, HumanMine includes genes, proteins, pathways, expression levels, SNPs, diseases and more, integrated into a single searchable database. HumanMine promotes integrative analysis, a powerful approach in modern biology that allows many sources of evidence to be analysed together. The data can be accessed through a user-friendly web interface as well as a powerful, scriptable web service API to allow programmatic access to data. The web interface includes a useful identifier resolution system, sophisticated query options and interactive results tables that enable powerful exploration of data, including data summaries, filtering, browsing and export. A set of graphical analysis tools provide a rich environment for data exploration including statistical enrichment of sets of genes or other biological entities. HumanMine can be used for integrative multi-staged analysis that can lead to new insights and uncover previously unknown relationships.

Database URL: https://www.humanmine.org

## Introduction

HumanMine (www.humanmine.org) is an integrated database of human genomics and proteomics data that provides a powerful interface to support sophisticated exploration and analysis of data compiled from experimental, computational and curated data sources. Built using the InterMine data integration platform, HumanMine includes genes, proteins, pathways, expression levels, SNPs, diseases and more, integrated into a single searchable database. Therefore HumanMine promotes integrative analysis, a powerful approach in modern biology that allows many sources of evidence to be analysed together. Data integration enables different pieces of information about a particular biological entity, or set of entities, to be surveyed together without the need to visit several databases. Such an approach has many advantages, including fewer false positives, compensation for missing or unreliable information and interrogation at different levels of genetic, genomic or proteomic regulation so helping in the understanding of complex biological systems. An important step in understanding such processes is also the interpretation of complementary data from model organisms. HumanMine interoperates with InterMine databases for a number of model organisms including mouse (MouseMine, (1)), rat (RatMine, (2)), zebrafish (ZebrafishMine, (3)), fly (FlyMine, (4)), worm (WormMine, http://intermine.wormbase.org/tools/wormmine), yeast (YeastMine,(4,5)) and Arabidopsis (ThaleMine, (6)). Here we describe how HumanMine can be used for integrative multi-staged analysis that can lead to new insights and uncover previously unknown relationships.

We will start by using the gene *PAX6* to illustrate the capabilities of HumanMine and then consider a more detailed use case in which HumanMine is used to support the exploration of disease comorbidities.

## InterMine

InterMine (7) is a data warehouse framework designed to facilitate the integration of diverse biological datasets and provide tools for bioinformatics analysis and visualisation both through a web interface and through extensive web services generated automatically from the underlying data model. InterMine was originally developed as FlyMine to address these issues for the *Drosophila* community. It has since been adopted by other model organism databases (MODs) including mouse (1), yeast (5), worm (http://intermine.wormbase.org), fish (3) and rat (2) and a growing number of databases covering other animals including, including Planaria (8), Hymenoptera (9), locusts (10) and armyworm (11). Plant species include Arabidopsis (6), Medicago (12), wheat (https://urgi.versailles.inra.fr/WheatMine), maize (11,13), and several from the legume family (14) (see registry.intermine.org). InterMine is also used for drug development (TargetMine (15)) and by large scale projects and consortia (PhytoMine (16), modMine (17), AllianceMine (https://www.alliancegenome.org/alliancemine)). HumanMine, described here, provides access to selected human data through the InterMine interface. A detailed description of the InterMine system is available elsewhere (7), but here we briefly review the data model and the relationships between different types of data. InterMine is able to integrate data from a wide variety of sources in various biological formats, including GFF3, FASTA, VCF, OBO, BioPAX (18), GAF, PSI (19) and Chado (20) and includes a powerful identifier resolution system such that any outdated identifiers from a dataset are mapped to the current ones. The underlying object-based core data model is based on the Sequence Ontology (SO (21)). This core is extended to include additional data types that do not fit to the SO, such as protein-protein interactions. Each type in the model (e.g. Gene, Disease) is called a class and is described through a set of attributes (such as gene symbol and gene length). References between classes link related data types (e.g. Gene and Allele) and facilitate navigation and exploration of the data. InterMine allows users to work with single items or lists of items (for example, a list of genes) and provides a number of search and analysis interfaces, including a keyword search, a genomic regions search and an advanced query builder, which exposes the data model and relationships and so allows the user to construct complex searches over the integrated data. In addition, a comprehensive web services API makes it straightforward to access data supplied by InterMine databases within programs, with support for most commonly used scripting languages (22). As many aspects of HumanMine conform to the FAIR principles of data management (23), the features described help to make analyses reproducible and the data more findable, accessible, interoperable and re-usable. HumanMine uses the recently released new interface to InterMine, BlueGenes (Manuscript in preparation).

## Data

HumanMine currently includes around 40 different data sets (supplementary table S1). The latest human reference genome and annotation release are loaded from the NCBI (https://www.ncbi.nlm.nih.gov/genome/?term=human[organism]). Data are focussed on high-throughput sets covering the genome, transcriptome, interactome and proteome and include full genome and proteome annotation, gene-pathway and Gene Ontology annotations, expression data including tissue- and disease-specific expression, proteome-wide and genetic interaction data, disease and phenotype data including Genome-Wide Association Studies, clinically significant SNPs and cis-eQTLs. A full list of data sources, including dates and versions, is available via the HumanMine home page (https://www.humanmine.org/humanmine/results/All%20data%20sources). Data are updated quarterly and we hope to include new data sources as they arise. The results reported here were generated using HumanMine version 12, released February 2022.

## HumanMine Access and help

HumanMine is freely available for the public to use, but creating an account allows a user to save custom queries and lists permanently. If a user is not logged in then lists and queries are saved for the duration of the session.

Extensive help documentation is available to aid in the use of both the user interface and the web services together with a number of video tutorials (http://intermine.org/intermine-user-docs/: http://intermine.org/im-docs/docs/web-services/index/#api-and-client-libraries).

## Exploratory data analysis: report pages and list analysis

The HumanMine interface provides a number of routes for exploring the integrated data, from keyword search and report pages to automated analysis of lists of data. For example, if a researcher wishes to explore data related to the *PAX6* gene they can use the simple keyword search. This returns links to genes, publications, interaction data, proteins, exons, UniProt protein features, mesh terms and a protein domain (figure 1). The keyword search accepts any identifier type, such as a gene symbol or ontology term and keywords and allows use of operators like AND, OR and NOT between terms. All data types within the database are searched and the results returned are faceted by class, making it easier to find the data of interest, for instance, the *PAX6* related proteins from Homo sapiens. Following the *Homo sapiens PAX6* gene link presents a report page collating much information about its function including expression, phenotypes, sequence variants, disease and homology (figure 2 - the report page has the following permanent URL, formed using the NCBI identifier for the gene, 5080: https://www.humanmine.org/humanmine/gene:5080). Specifically, we learn that *PAX6* is expressed in the developing eye and brain, is involved in transcription and that it is associated with a number of eye-related disorders such as aniridia, keratitis and Peters Anomaly. Examination of gene expression and protein localisation data shows us that *PAX6* is expressed not only in various brain structures but also in the pancreas and is upregulated in various cancers (figure 2).

**Figure1:**
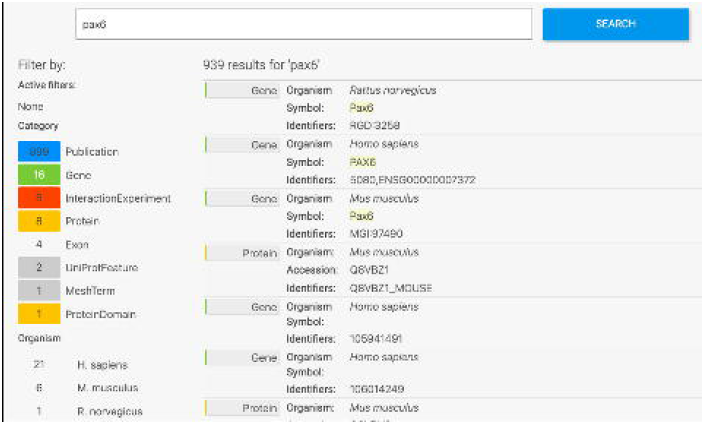
Keyword search. A search for “Pax6” returns a filterable menu of data classes and organisms with the number of entities found in each on the left and individual entities displayed on the right. Each individual entity provides a link to its report page.

**Figure2:**
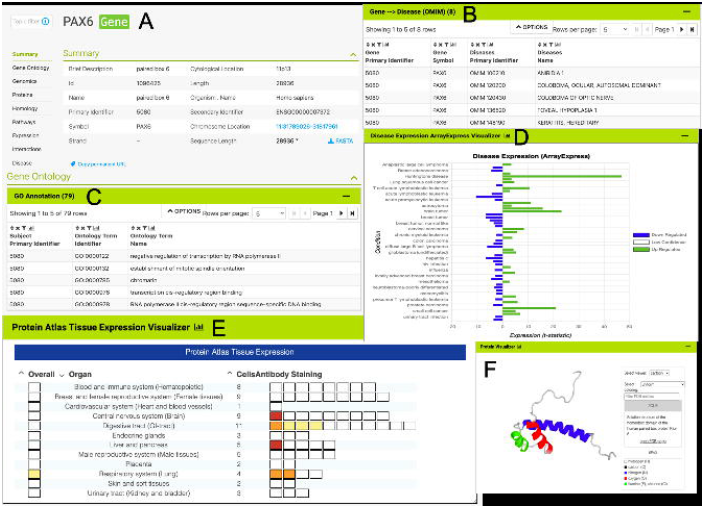
The *PAX6* report page. Report pages present data through a range of interactive tables, graphs and visualisations depending on the data type. A selection of features from the report page for the human *PAX6* gene are shown here. A) A summary of the main identifiers and chromosomal location. B). An interactive table of Gene Ontology annotations. Only the first 5 rows are shown. C). A table of disease annotations (original data source: OMIM, https://www.omim.org). Only the first 5 rows are displayed. D). A graph showing up and down-regulation of the *PAX6* gene in various disease conditions (original data from ArrayExpress experiment E-MTAB-62, https://www.ebi.ac.uk/arrayexpress/experiments/E-MTAB-62). E). A graph showing protein localisation data (from the Protein Atlas project, https://www.proteinatlas.org/humanproteome/tissue.). F). A protein structure viewer pulling in data from the Protein Data Bank (https://www.rcsb.org).

Within the report page for the *PAX6* gene, it is also possible to identify interesting gene sets (“lists”) in which *PAX6* is found. For instance, *PAX6* is found in a number of Genomics England gene panels, highlighting its importance in many areas of development. Selecting the list “PL_GenomicsEngland_GenePanel:Glaucoma_(developmental)”, from the *PAX6* report page, allows us to explore aggregated data for this set of genes. Alongside a table detailing the individual members of the list, enrichment statistics show Gene Ontology terms for various visual and sensory organ development (such as iris morphogenesis (p-value: 1.20e-3) and eye development (p-value: 1.57e-2) while enriched pathway terms include TRAF3-dependent IRF activation pathway (p-value: 7.83e-4). Widgets and tools then allow us to examine the chromosome distribution of the genes in the list, identify interacting partners through both a table and a network graph, examine heat maps of gene expression, visualise pathway interactions and Gene-GO term relationships (figure 3).

**Figure 3:**
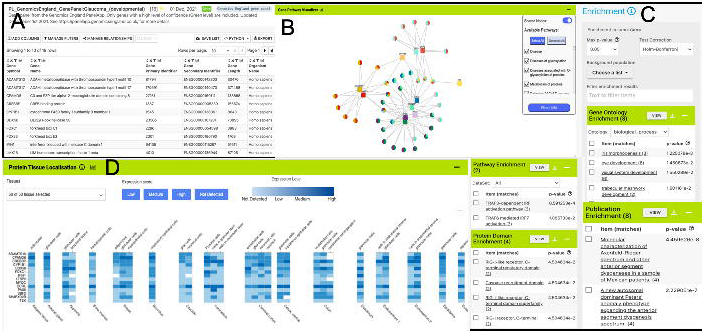
List analysis page for the public list: PL_GenomicsEngland_GenePanel:Glaucoma_(developmental). Like the report pages, list analysis pages provide a number of interactive tables and graphs. A selection is shown here. A). Interactive table summarising the contents of the list. B) A network graph showing Gene-Pathway connections for genes in the list. Only genes that have 2 or more pathway connections are shown (this option can be toggled on the menu panel). The menu panel also allows filtering of the pathway annotations used in the graph. C) Enrichment statistics for Gene Ontology terms, Publications, Protein domains and Pathway annotations. D) A heat map showing protein localisation for each gene in the list (original data from The Protein Atlas project, https://www.proteinatlas.org/humanproteome/tissue).

The ability to create and use lists of entities is integral to data analysis within HumanMine. All lists can be viewed with such list analysis pages which, like the report pages, provide summarised data for the contents of the list. Lists of items with standardised annotations, such as Gene Ontology annotations, facilitate analysis by statistical enrichment methods. InterMine uses the hypergeometric distribution with a choice of multiple test corrections to provide enrichment statistics for lists of genes i.e. to discover whether an annotation is unexpectedly frequent within the group. Interactive tables of enrichment p-values, calculated for Gene Ontology terms, pathway annotations, protein domain annotations and publications are displayed on the gene list analysis pages. The tables can be adjusted for test correction, p-value threshold and background reference set (the default being all genes in the human genome with an annotation of the type being analysed, e.g. all human genes that have associated GO annotations) and subsets of the genes with a particular enriched term can easily be viewed and saved as further lists. In addition to the enrichment tables, a tools library allows for the easy addition of visualisations to the list pages by the database maintainer.

Lists created within HumanMine or from an external source can be saved into the user’s account for further use. In addition, HumanMine provides a number of public lists (sets of interesting genes from relevant publications or resources) such as the sets of rare-disease and cancer genes defined by the Genomics England Gene PanelApp project (https://panelapp.genomicsengland.co.uk, (24)) (only “Green” level panels are included, which have a high level of evidence for the gene-disease association). Such public lists have a number of applications including identifying whether a gene of interest is present in any of the lists (shown on gene report pages), notifying users of significant overlaps between their lists and the public lists (currently available through the web services but soon to be developed for the user interface) and simply as a convenient list of genes known to be significantly involved in a particular disease.

All lists, both public and belonging to a user, can be viewed on the “Lists” page, which has a number of functions for the organisation and exploration of the lists available, including the addition of free-text descriptive annotation as “tags”, creation of folders, as well as search and filtering functions (figure 4). Two or more lists can be combined using set operations: union, intersection, difference and subtraction.

**Figure 4:**
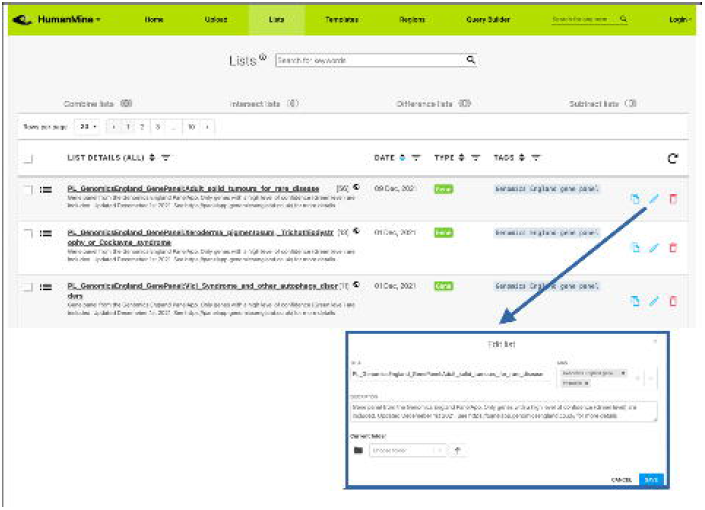
Lists. All lists, both public and private for a user account, can be viewed under the Lists tab.

## Structured Data Analysis: template searches and the query builder

After exploring the report pages for a particular entity, a researcher may wish to ask more structured or focussed questions about the data. Continuing with our *PAX6* example, we noted earlier that *PAX6* is upregulated in a number of cancers, including small cell lung cancer. To investigate this further a researcher may follow a number of lines of investigation. For example, to 1) identify other genes upregulated in small cell lung cancer and 2) identify whether any of these genes interact with *PAX6.* A library of ‘canned’ or pre-formed searches, called template searches, allow the user to easily search across any integrated data set (figure 5). For example, the above two questions about *PAX6* could be investigated with a single template search, “Gene(s) + Disease --> Interactors + Disease Expression”. This template combines disease expression data (ArrayExpress experiment E-MTAB-62 (25)) and interaction data (IntAct (26) and BioGrid (27)) and allows us to search for genes that interact with *PAX6* that are, according to the data, also up-regulated in a defined disease state, here lung cancer.

**Figure 5:**
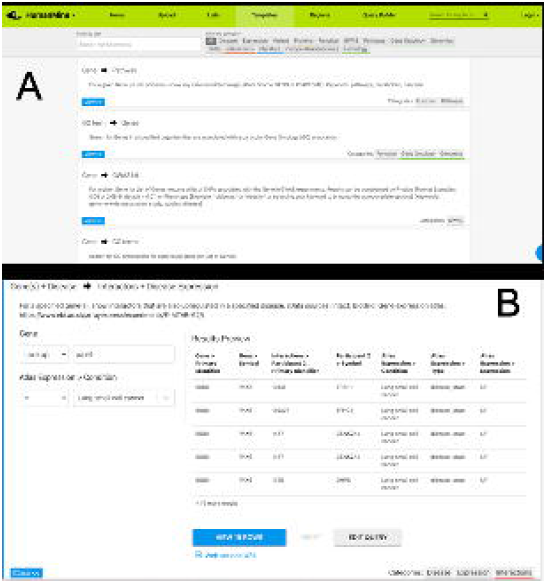
A) A library of “template” searches is available from the “Templates” tab. The template library can be searched using keywords or filtered using the various data category tags. B) The “Gene(s) + Disease Interactors + Disease Expression” template expanded. Each template provides one or more constraints that can be modified according to the search the user wishes to run. A preview of the template result is shown with options to view the full results or edit the query in the Query Builder.

A template search typically has one or more *start* data types as input and one or more *end* data types that are provided as output and allows the user to add constraints (or filters) to these to control the subset of data returned. Templates cover both simple searches and more complex searches combining two or more datasets, and in most cases can be executed with either a single item (e.g. a gene) or a list of items (such as a pre-saved gene list).

More advanced data mining is available through the query builder, which can be used to generate *de novo* queries or to modify template searches. For instance, the template “Gene(s) + Disease --> Interactors + Disease Expression” can be displayed in the query builder and can be modified to return genes that interact with *PAX6* that are expressed in lung squamous cell cancer as well as small cell lung cancer (figure 6). The query builder interface displays a hierarchical data model browser through which classes of data can be selected for combining into a search. Constraints that filter the data and the results that are provided as output can be defined through the interface together with more advanced features such as constraint logic (data types are combined with an “AND” by default but can be configured for OR, see figure 6) and data join type, i.e. whether the intersection of two datasets should be returned or the union of the datasets (the latter being returned by default). For instance, in our example, it is possible to return either all interacting genes regardless of whether they also show expression in the defined conditions, or only return those interactors for which the expression conditions are satisfied. Template searches and the Query Builder present results as InterMine Results Tables. From here, we can, for example, use a column summary function to find that 1) 18 interacting genes are returned and 2) that, using the column summary to filter the table, 14 of the genes are associated with small cell lung cancer and 12 with lung squamous cell cancer (figure 7). The “Save as list” function of the results table allows us to save the set of 14 upregulated small cell lung cancer genes that interact with *PAX6* enabling this set to be further analysed through the list analysis page, further template searches and set operations on lists. The Gene Pathway Visualizer on the list analysis page, for example, highlights a number of potentially important cancer genes including SOX2, CSNK2A1, LMX2, SMAD5, APP and TP73 (figure 8). The intersection of these genes with a set known to be involved in small cell lung cancer from the DisGenNet (28) dataset confirms that two, SOX2 and TP73 are known to be directly implicated in small cell lung cancers, suggesting that the rest of the list may also be worthy of further investigation in the context of lung cancer.

**Figure 6:**
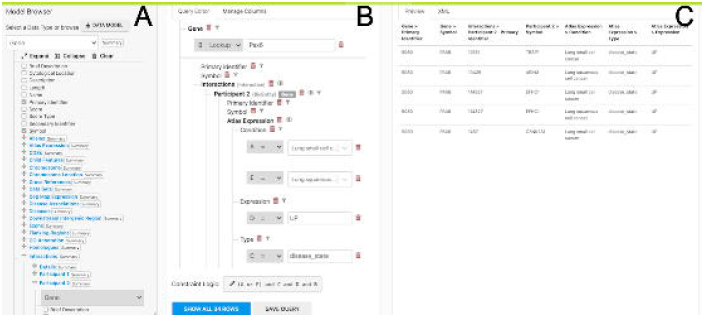
The Query Builder. The query builder consists of three main panels - the model Browser (A), the query editor (B) and the query preview (C). The “Gene(s) + Disease Interactors + Disease Expression” search is shown with an extra constraint for “condition = Lung squamous cell cancer” added. Each constraint in the query editor is labelled with a letter enabling the constraint logic to be edited here to give ((A or E) and C and D and B).

**Figure 7:**
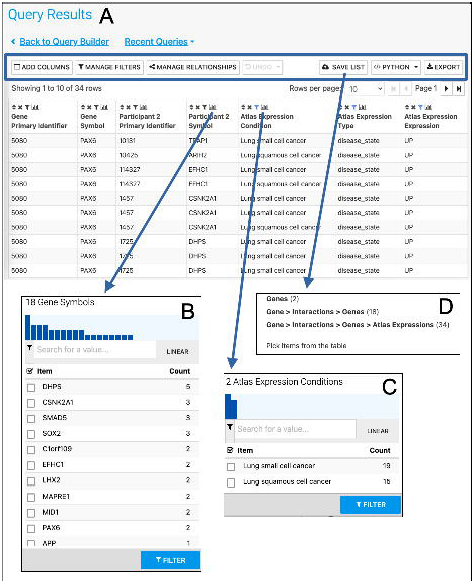
The Results Table showing results from the query “Gene(s) + Disease Interactors + Disease Expression” for the *PAX6* gene with constraints on the disease name for Small cell lung cancer or Lung squamous cell cancer. The results tables provide many additional functions including “Add columns” allowing additional data to be added, “Manage filters” allowing filters on any column to be defined, “Manage relationships” enabling either the union or intersect of classes of related data in the table to be defined, “Save list” enabling subsets of items in the table to be saved as lists, “Python”, automatic code generation, available as a drop-down list of available languages and “Export” (A). The column summary on the Participant 2 > Symbol column (the genes with which *PAX6* interacts) allows the number of unique interacting genes (18) to be found (B). Using the column summary on the Atlas Expression > Condition column it is possible to see the number of rows for each disease condition. This could be used to filter the table to show just one of the disease conditions (C). The “Save as list” function can be used to save any set of items from the table. Here it may be useful to save the set of interacting genes (Gene > Interactions > Genes (18)) (D). To save the set of interacting genes specific to one of the cancers, the table could be filtered first using the column summary function as described above.

**Figure 8:**
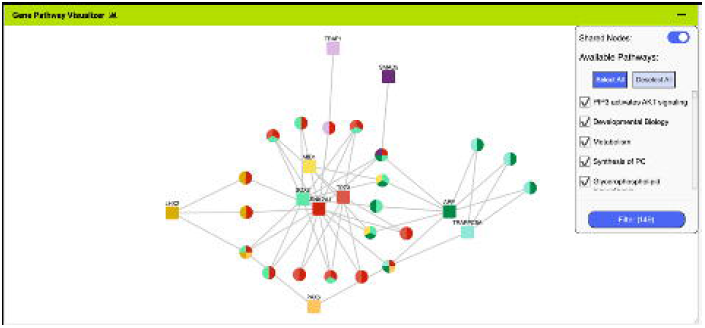
The Gene Pathway Visualizer on the list analysis page, showing Gene - Pathway associations for the 14 upregulated small cell lung cancer genes that interact with *PAX6*. A number of potentially important cancer genes including SOX2, CSNK2A1, LMX2, SMAD5, APP and TP73 can be seen. Only genes that have 2 or more pathway connections are shown (this option can be toggled on the menu panel).

Thus the Results Table is a powerful query analysis tool in itself, providing many features for the manipulation of the results, including the column summaries (simple statistics describing the data in a column including the number of distinct attributes for categorical data, or, for numerical data, the mean, median, minimum and maximum), data filtering, sorting, list generation and the ability to add additional columns of data (figure 7). The results tables can be exported in common formats including TAB/CSV and as a frictionless data package (https://frictionlessdata.io). Nucleotide and protein sequences can be exported in FASTA format. An additional feature of the results tables in the new BlueGenes interface is the automatic generation of enrichment statistics and the inclusion of analysis tools as seen on the list analysis pages (figure 3). The enrichment is based on the first column of genes or proteins, however, if more than one column of genes or proteins is present in the table, another column can be selected for the enrichment calculations from the settings bar.

## Using HumanMine to explore external data

We learnt from the GO annotations on the report page that *PAX6* is a transcription factor involved in many cellular functions. It follows, therefore, that a researcher may wish to analyse genes that are regulated by *PAX6.* HumanMine provides two interfaces for the upload of external data, a list upload and a region search. For instance, a set of human genes already identified as *PAX6* target genes (29) can be uploaded to HumanMine (HumanMine public list PL_Pax6_Targets) using the list upload function enabling them to be analysed alongside the integrated data using any of the functionality discussed above. The template “Gene → Gene Expression (Tissue, Disease; Array Express, E-MTAB-62)” can be run using this list as input and with “Atlas Expression > type” set to disease_state and “Condition” set to “Contains” Lung. The resulting results table can be further filtered using the Atlas Expression Condition column summary to show data for just “small cell lung cancer” and “Lung small cell cancer” so identifying 51 upregulated genes, five of which (ASCL1, EPHA7, ID2, RUNXITI and SOX2) already have a known involvement in small cell lung cancer according to the DisGenNet data, with the others therefore providing a set of candidate genes for further research.

HumanMine allows users to upload any lists of objects for which data classes exist (for instance genes, proteins, transcripts). The list upload system includes a powerful identifier resolution system such that any outdated identifiers or synonyms can be updated to the current human annotation. For genes, HumanMine supports NCBI Entrez (30) gene identifiers, Ensembl (31) gene identifiers and HGNC (32) identifiers, with the main genome annotation being from NCBI. Thus, once a list has been uploaded it can be analysed in the same way as a list generated from within the database and will benefit from an automatically generated list analysis page.

The Region Search tool makes it possible to identify, for specified genomic spans, genomic sequence features such as genes and regulatory regions that overlap the spans. Thus the list of *PAX6* target genes could equally have been created by using a published set of ChIP-seq binding intervals (29) and searching for overlapping genes. The region search results interface allows such a set of genes to be easily saved as a list. The genomic regions can be extended by specified amounts beyond the 5’ and 3’ coordinates given, and a strand-specific search can be carried out if required. Features identified can be saved as lists or exported, either individually for each region specified or for all regions searched.

## Web services

In addition to the user interface, HumanMine can be accessed through a comprehensive web services layer (12) through which all the functionality of the user interface can be accessed. Client libraries in several languages are available (http://intermine.readthedocs.io/en/latest/web-services/#api-and-client-libraries) or RESTful endpoints may be accessed directly (http://iodocs.apps.intermine.org/). Automatic code generation through the user interface provides an easy starting point for those wishing to use the web services. For example, code for running the template mentioned earlier, “Gene → Gene Expression (Tissue, Disease; Array Express, E-MTAB-62)”, can be generated directly from the template form (figure 9). This code can then be added to a script that has imported the corresponding client library, so facilitating script-based analysis in which data from InterMine is dynamically retrieved. In addition, an R library, InteMineR (33) available from BioConductor (https://www.bioconductor.org/packages/release/bioc/html/InterMineR.html), allows queries against HumanMine to be run using the R programming language.

**Figure 9:**
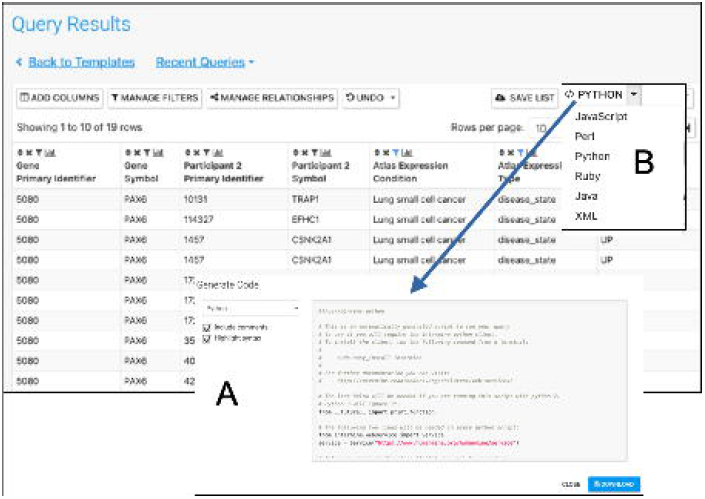
Automatic code generation. From any results table, it is possible to view and copy code for the underlying query in various programming languages. Here the python code for the Gene(s) + Disease Interactors + Disease Expression result is shown (A). Code for Python, Perl, Ruby, Javascript and Java is available (B).

## Use-Case: using HumanMine to explore shared pathways in disease comorbidities

The combination of integrated data and analysis tools in HumanMine allows for the exploration of shared genes and pathways between two or more conditions. Asthma and type 2 diabetes are two of the most prevalent chronic diseases, presenting significant challenges to public health. While patients frequently only have one of either asthma or type 2 diabetes, the two diseases also have a well-documented but poorly understood comorbidity, in which they co-occur more frequently than would be expected by chance (34). Studying disease comorbidities allows both a better understanding of the pathology of the respective diseases, as well as identification of the underlying mechanisms of the comorbidity, which may provide opportunities for more targeted interventions and therapies.

For this example, data from multiple disease sources (OMIM (35), ClinVar (36), GWAS Catalogue (37), DisGenNet (28) and the Human Phenotype Ontology, HPO (38)) were searched for genes associated with either Type 2 diabetes or Asthma. The resulting lists of genes for each disease were combined by using the list intersection function to identify genes shared between the two diseases. Analysis of the resulting shared set of 912 genes (from now on called the shared set, supplementary table S2) provides a number of insights into the underlying causes of the comorbidity, some of which are discussed below (figure 10). The enriched Gene Ontology terms and Pathway annotations for the shared set identify a number of underlying shared biological functions between the two conditions, in particular in the immune system, signal transduction and hemostasis (supplementary table S3 A and B).

**Figure 10:**
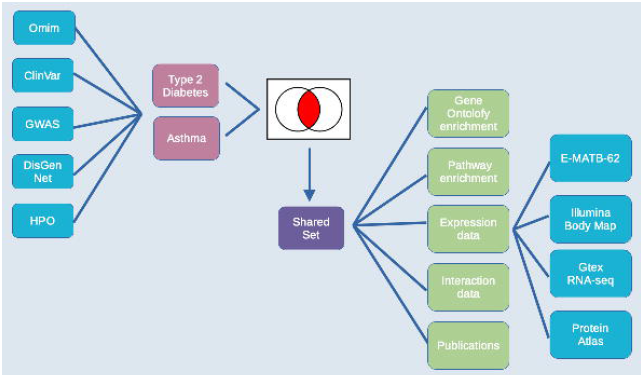
Exploring comorbidities using HumanMine. A schematic representation of the steps involved in the use-case “Using HumanMine to explore shared pathways in disease comorbidities”. See text for details.

There is evidence to suggest that asthma may increase the risk of developing type 2 diabetes (39). The mechanisms by which this increased risk may occur, however, are not fully understood. It has been suggested that increased levels of some inflammatory cytokines known to be present in asthma may contribute to the development of type 2 diabetes (39). This proposal is backed up here by the finding that 249 genes from the “shared set” are annotated with the high-level GO term “Inflammatory response” and the enrichment of a number of pathways involved in inflammation, particularly the interleukin signalling pathways. The proinflammatory biomarkers interleukin-6 (IL6), C-reactive protein (CRP) and TNF-alpha (TNF), present in the “shared set” and known to be elevated in low-grade inflammation, are thought to be major contributors to the development of type 2 diabetes (40), (41). Thus, it is possible that increased circulating levels of some inflammatory cytokines known to be present in asthma may also contribute to the development of type 2 diabetes.

Another important immune gene, TGFB1, present in the shared list, encodes the cytokine TGF-beta 1. Its roles in airway inflammation and remodelling in asthma are well-defined (42), and asthma severity is associated with different TGFB1 polymorphisms in humans (43). The role of TGF-beta 1 in type 2 diabetes is less established, however, an interesting hypothesis is that it drives dedifferentiation of beta cells (44), a process implicated in type 2 diabetes pathogenesis (45): dedifferentiation is also part of asthmatic airway remodelling (46). Targeting TGF-beta 1 remains challenging (47) but could offer new therapeutic potential for both diseases.

Toll-like receptors (TLRs) are expressed on the membranes of leukocytes and activate immune cell responses in response to microbes. Such activation of TLRs in the lung can induce inflammation, inflammatory cell recruitment and cytokine release. In particular, TLR2 and TLR4 have been shown to sustain the inflammatory response in asthma (48). There is also evidence that TLR4 is directly involved in the pathophysiology of type 2 diabetes (49), (49,50) and it has been suggested that drugs targeting the TLR4 signal pathway could be candidates for the treatment of inflammatory co-morbidities (51). The “Pathway → Genes” template was used to retrieve all known genes of the TLR4 signalling pathway. List intersection shows that 45 genes from the shared set function in this pathway (out of a total of 128 pathway genes). Interestingly 34 of these 45 proteins have a drug entry in DrugBank (52) [Source: TargetMine (15): https://targetmine.mizuguchilab.org], reinforcing the therapeutic potential of this pathway.

A number of other genes in the shared set warrant further analysis. For example, the enriched pathways include a number related to GPCR signalling with 128 genes being annotated with “signalling by GPCR”. Included in these are BDKRB1 and BDKRB2, receptors for pro-inflammatory bradykinins, which have an established role in asthma (53) but less is known about their roles in type 2 diabetes. BDKRB1 has previously been identified as upregulated in allergic airway inflammation (54) and a mouse model of diabetes (55) and maybe worth further investigation.

The HumanMine integrated data allows many other datasets to be examined alongside the disease data. For instance, five datasets allow investigation of gene expression at the tissue level (E-MTAB-62 and E-MTAB-513 (from https://www.ebi.ac.uk/arrayexpress), GTex (from https://gtexportal.org) and Human Protein Atlas (both protein localisation data and RNA-seq data, (56))). Such data allow other factors to be considered in the above analysis, for instance, genes from the shared set that are expressed in both the lung and adipose tissue. An initial analysis using the Human Protein Atlas (56) RNA-seq data indicates that 406 of the genes from the shared set are expressed in both tissues. These are enriched for immune system and signal transduction pathways as discussed above and include IL6, TLR2 and TLR4. The expression viewers on the list analysis page allow for a visual overview of the genes in the shared list that are expressed in the lung and adipose tissue (figure 11). Each expression set provides a slightly different result, emphasising the benefit of being able to examine multiple datasets alongside each other.

**Figure 11:**
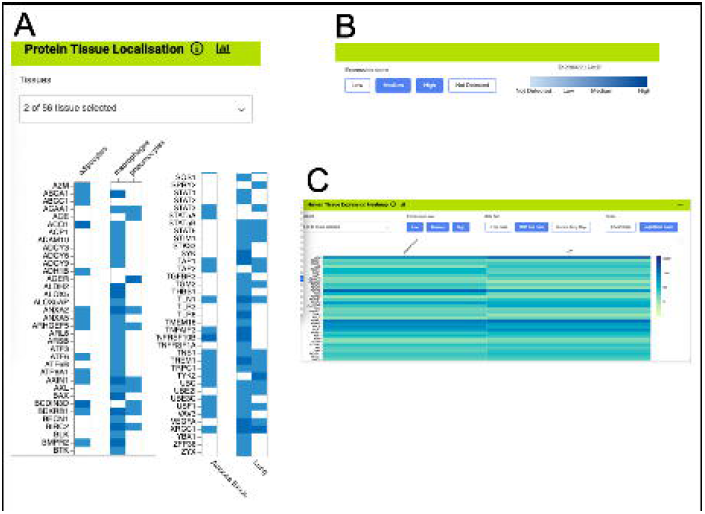
Gene expression heat maps for a selection of the genes from the shared set. A). Protein tissue localisation (original data from The Protein Atlas project, https://www.proteinatlas.org/humanproteome/tissue). The viewer has been filtered to show data for adipose tissue (adipocytes) and lung (macrophages and pneumocytes) only. B). It is possible to toggle the expression score ‘bins’ on or off and a colour scale representing expression level is shown. C. Heat map filtered to show RNA-seq data for adipose and lung (original data from The Protein Atlas project, https://www.proteinatlas.org/humanproteome/tissue). The viewer allows toggling between other expression data sets, showing different binned levels of expression and provides a scale for expression level.

Expanding the analysis further, the network of interacting proteins for the shared set can be examined, the interacting partners of each gene being found using the “gene → interactions” template. Enrichment analysis of the resulting “expanded” set highlights signalling by Rho GTPases. The RHO GTPases are emerging regulators of glucose homeostasis and have been implicated in beta cell dysfunction (57). They are also well known to play an important role in the pathophysiology of asthma and recent advances have suggested novel roles for RhoA in regulating allergic airway inflammation. The RHO GTPases are promising targets for therapeutic treatment of asthma and two major Rho-kinase inhibitors have been developed (58).

A further way to enhance the analysis would be to investigate the literature for publications exploring the asthma/diabetes comorbidity, for instance by searching for papers that mention both in either their titles or abstracts. In addition to searching data, HumanMine makes it easy to begin a literature search for the interesting sets of genes identified, by means of the PubMed-to-gene mappings provided by PubMed. Editing the template “Gene → Publications” in the query builder to remove the gene constraint and add two constraints on the publication title (figure 12) allows publications in which the title includes both Asthma and Diabetes to be easily identified. Seven publications are returned by the search, interestingly identifying a TLR2 polymorphism associated with type 1 diabetes and allergic asthma (figure 12). In addition, the publication enrichment for the shared set can also be searched for interesting clues. Although no papers directly analysing diabetes and asthma appear to cite an unexpectedly high number of the shared set members, there are some potentially interesting leads - for instance, 37 genes from a paper entitled “Analyses of shared genetic factors between asthma and obesity in children”.

**Figure 12:**
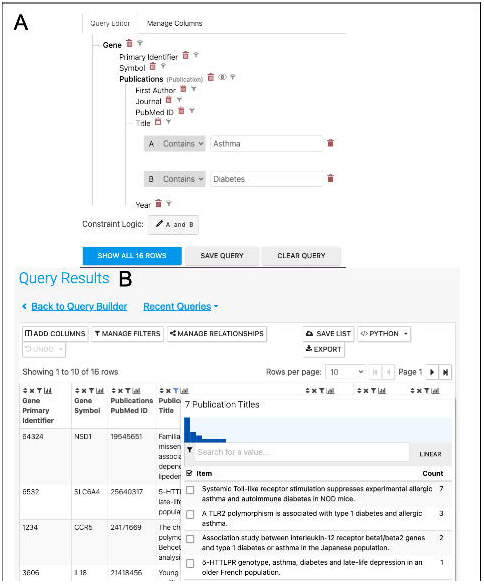
A). A query to find publications in which the title includes both Asthma and Diabetes. B). The results return seven publications as shown by the column summary.

While it is not possible to provide a full analysis of the data and its implications in therapeutic treatment of asthma and diabetes here, the above use-case illustrates how HumanMine facilitates an in-depth exploration of the data available and highlights many areas where further research could be focussed. The method outlined above can be applied to any combination of disease comorbidity and can easily be adapted to other research areas.

## Conclusion

HumanMine is an advanced biological data warehouse for human genomics and proteomics data, exposing the data through intuitive graphical user interfaces and as well as allowing programs to access data through extensive web services. The *PAX6* and disease comorbidity examples illustrate how HumanMine can be used for powerful explorative and iterative data analysis. The combination of list creation, list operations, querying and filtering allows a workflow of tasks to be built up, the results of which can be examined at various stages using enrichment tools, list tools and report pages (figure 13). Furthermore, the combination of automatic code generation and web services makes it easy to replicate such steps in a script and so repeat the analysis at scale with different starting conditions and parameters.

**Figure 13:**
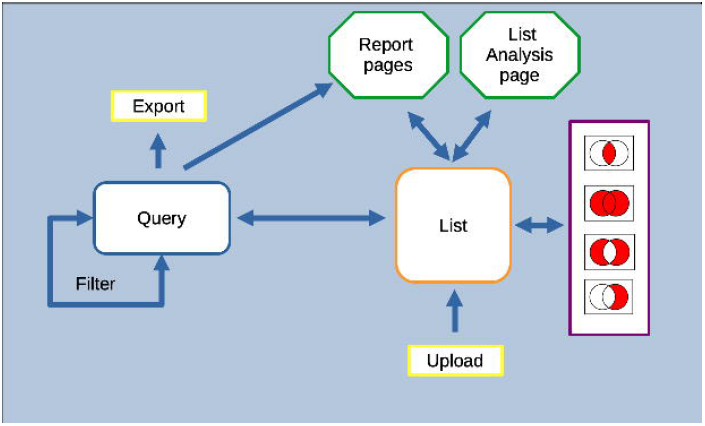
Features of the InterMine interface can be combined to create iterative workflows. For instance, the entities from the results of a query generated either using the query builder or through a template search can be saved as a list. The list can be fed into further queries, or combined with other lists using set operations to create further lists. At any stage, individual entities or lists can be examined in more detail through the list analysis and report pages.

---

Supplementary Table S1: The key data sources present in the HumanMine database v12.

Supplementary Table S2: The “Shared set” of genes. The set of genes found to be associated with both Asthma and Type 2 diabetes.

Supplementary Table S3A: The Gene Ontology term enrichment for the shared set of genes. The P-value is calculated using the hypergeometric distribution, with the background being all human genes that have a Gene Ontology annotation.

Supplementary Table S3B: Pathway term enrichment for the shared set of genes. The P-value is calculated using the hypergeometric distribution, with the background being all human genes with a Pathway annotation. The pathway data originate from the Reactome database (59)

